# Efficient and scalable *de novo* protein design using a relaxed sequence space

**DOI:** 10.1101/2023.02.24.529906

**Authors:** Christopher Frank, Ali Khoshouei, Yosta de Stigter, Dominik Schiewitz, Shihao Feng, Sergey Ovchinnikov, Hendrik Dietz

## Abstract

Deep learning techniques are being used to design new proteins by creating target backbone geometries and finding sequences that can fold into those shapes. While methods like ProteinMPNN provide an efficient algorithm for generating sequences for a given protein backbone, there is still room for improving the scope and computational efficiency of backbone generation. Here, we report a backbone hallucination protocol that uses a relaxed sequence representation. Our method enables protein backbone generation using a gradient descent driven hallucination approach and offers orders-of-magnitude efficiency enhancements over previous hallucination approaches. We designed and experimentally produced over 50 proteins, most of which expressed well in E. Coli, were soluble and adopted the desired oligomeric state along with the correct composition of secondary structure as measured by CD. Exemplarily, *we* determined 3D electron density maps using single-particle cryo EM analysis for three single-chain *de-novo* proteins comprising 600 AA which closely matched with the designed shape. These have no structural analogues in the protein data bank (PDB), representing potentially novel folds or arrangement of domains. Our approach broadens the scope of de novo protein design and contributes to accessibility to a wider community.

## Introduction

Deep-Learning (DL) based protein structure prediction methods such as AlphaFold2 (AF2)^1^ and RoseTTAFold^2^ can generalize beyond the sequence and structure space they have been trained upon to correctly predict the structures of *de novo* designed proteins^3–6^. Additionally, AlphaFold2 can distinguish between favorable and unfavorable structure templates^7^, suggesting that it has learned the physical properties of energetically stable protein backbones. DL-based structure prediction methods can thus connect sequence space and protein structure space, and enable searching for protein structures that satisfy a given design target. A DL based design method called ‘deep network hallucination’^8^ leverages this connection for a variety of protein design problems by iteratively updating a protein sequence until a desired property encoded in a mathematical loss function is obtained. Typically, when performing deep network hallucination, random mutations are applied in a Monte Carlo Markov Chain (MCMC) fashion^4,6,9–11^. However, this method can be computationally inefficient due to the need for a large number of iterations. This approach may also fail to identify a solution, similar to issues faced by numerical solutions to mathematical problems in a random Monte Carlo fashion. To improve the likelihood of generating accurate predictions, researchers have proposed using sequence gradients obtained by inverting structure prediction networks^12^. However, when updating the gradients, a major issue arises due to the discrete, one-hot encoded representation of sequences. One-hot encoding (Fig.: 1, A) represents each amino acid or nucleotide as a binary vector with all zeros, except for a single ‘1’, indicating the position of the corresponding residue. While the obtained gradient from the backpropagation through the structure prediction network is non discrete, previous approaches tried to regain discreteness by a ‘straight-through estimator’^13^ in the sequence update loop^5,12,14^. Though this approach worked well for models predicting distribution of distances for every pair of positions, such as TrRosetta, the approach did not work well in more recent models such as AlphaFold, where single amino acid changes could radically alter the predicted structure, resulting in unstable and inefficient optimization. Therefore, alternative methods of representing sequences in a continuous format are being explored to overcome this problem.

**Figure 1.**
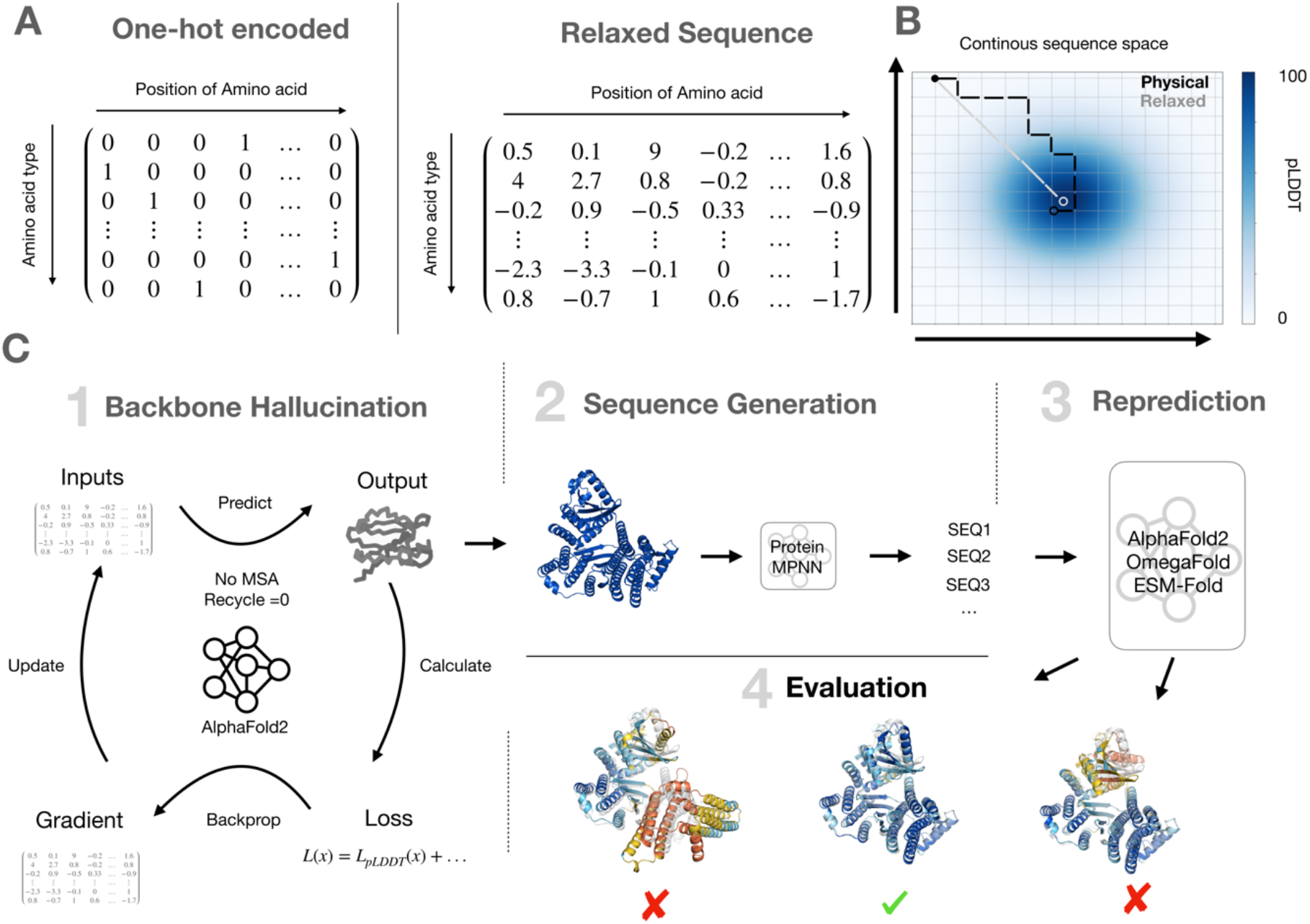
Schematics of the protein backbone design pipeline. A) Schematic representation of the difference between a physical input sequence that is one hot encoded and a relaxed sequence that uses float numbers. B) Schematic representation of the continuous sequence space used for relaxed sequence hallucination. Grid lines indicate physical sequences, while the color shows the pLDDT of the sequences. The grey line shows the restriction free GD process used for relaxed sequence hallucination, while the black line shows an exemplary trajectory when doing GD with physical sequences. C) Schematic representation of the relaxed sequence design pipeline consisting of: 1) Backbone hallucination 2) sequence generation with Protein MPNN 3) Reprediction with AF2, EMS-Fold and OmegaFold 4) evaluation of predictions.

The AF2 network can predict sequences generated through the hallucination process with high confidence. However, despite this high confidence, the experimental success rate of the resulting proteins is often low, resulting in the production of insoluble proteins^6,15^. To resolve this issue, it has become standard practice to generate a new sequence for the given backbone using the ProteinMPNN sequence design software^15^. ProteinMPNN has demonstrated significant success in designing soluble sequences for specific protein backbones, making it a valuable tool for improving the experimental success rate of the resulting proteins. Since the sequence design process for a given backbone geometry can be reliably handled with ProteinMPNN, we hypothesized that using a continuous (relaxed) sequence representation would allow for a more targeted and efficient gradient-descent search through the structure space for desired backbone properties while avoiding the issues associated with the need for a discrete, one-hot encoded sequence. A relaxed sequence representation approach may thus allow for more precise and effective exploration of the structure space for backbones, resulting in improved outcomes for protein design.

To test our hypothesis, we utilized a representation where each amino acid position in the input sequence is a vector of 20 values representing the contributions from each of the 20 natural amino acids. These values can be fractional, greater than one, or negative and positive in value (Fig.: 1, B). Though the input sequence, termed “logits”, may represent an unnormalized probability distribution of amino acids, in this work they serve as a means to more efficiently explore the AF2 network for backbone geometries. Specifically, the relaxed sequence representation enables adopting a gradient-descent based hallucination protocol that optimizes a loss function with rapid convergence towards high confidence protein structures (Fig.: 1, C). To implement this relaxed sequence representation, we used the ColabDesign framework, which offers various deep network hallucination protocols with a modified, differentiable version of AF2.

Using this approach for backbone generation, we set up a protein design pipeline that uses relaxed sequence hallucination to generate target backbone geometries. After obtaining high-confidence backbone coordinates with a pLDDT score greater than 90, we then use the ProteinMPNN network to generate actual physical candidate sequences represented as a one-hot encoding (Fig.: 1, D). We then repredict the candidate sequences by AlphaFold2 and ESM-fold^16^ to validate the one-hot candidate sequences obtained from ProteinMPNN with respect to forming the input backbone geometry. This step is necessary since AF2 models P(structure | sequence) whereas MPNN models P(sequence | structure). It is thus possible that P(sequence | wrong_structure) >= P(sequence | desired_structure). Therefore one needs to confirm that P(desired_structure | sequence) is valid using a new pass through a structure prediction model.

### Computational design pipeline & benchmarking

To test the efficiency of our relaxed sequence hallucination, we performed a benchmarking experiment in which we unconditionally hallucinated backbones with three different protocols: gradient-descent with relaxed sequence (GD relaxed); GD with argmax() and one-hot encoded sequences (GD hard); and Markov-Chain-Monte-Carlo search (MCMC) within the ColabDesign framework (Fig.: 2, A). GD relaxed robustly outperformed the other design approaches in all cases tested: the average convergence “half-life” was 20 iterations across all design test cases, whereas GD hard and MCMC failed to converge to satisfactory results (pLDDT >90) within the tested number of sequence update steps (Fig.: 2, A & Supp. Fig.: 1, A). Notably the GD relaxed method reliably produced high confidence backbones with pLDDT score larger than 95 for all unconditional hallucination trajectories tested (Fig.: 2, A). We note GD relaxed utilizes a back propagation step to implement the gradient descent, as opposed to only performing forward passes through AF2 as in MCMC. Hence, the computational effort for one sequence update step is greater in GD relaxed, but because of the orders-of-magnitude reduction in steps needed to convergence, relaxed sequence hallucination strongly outperforms MCMC in terms of design speed (Supp. Fig.: 1, B).

**Figure 2.**
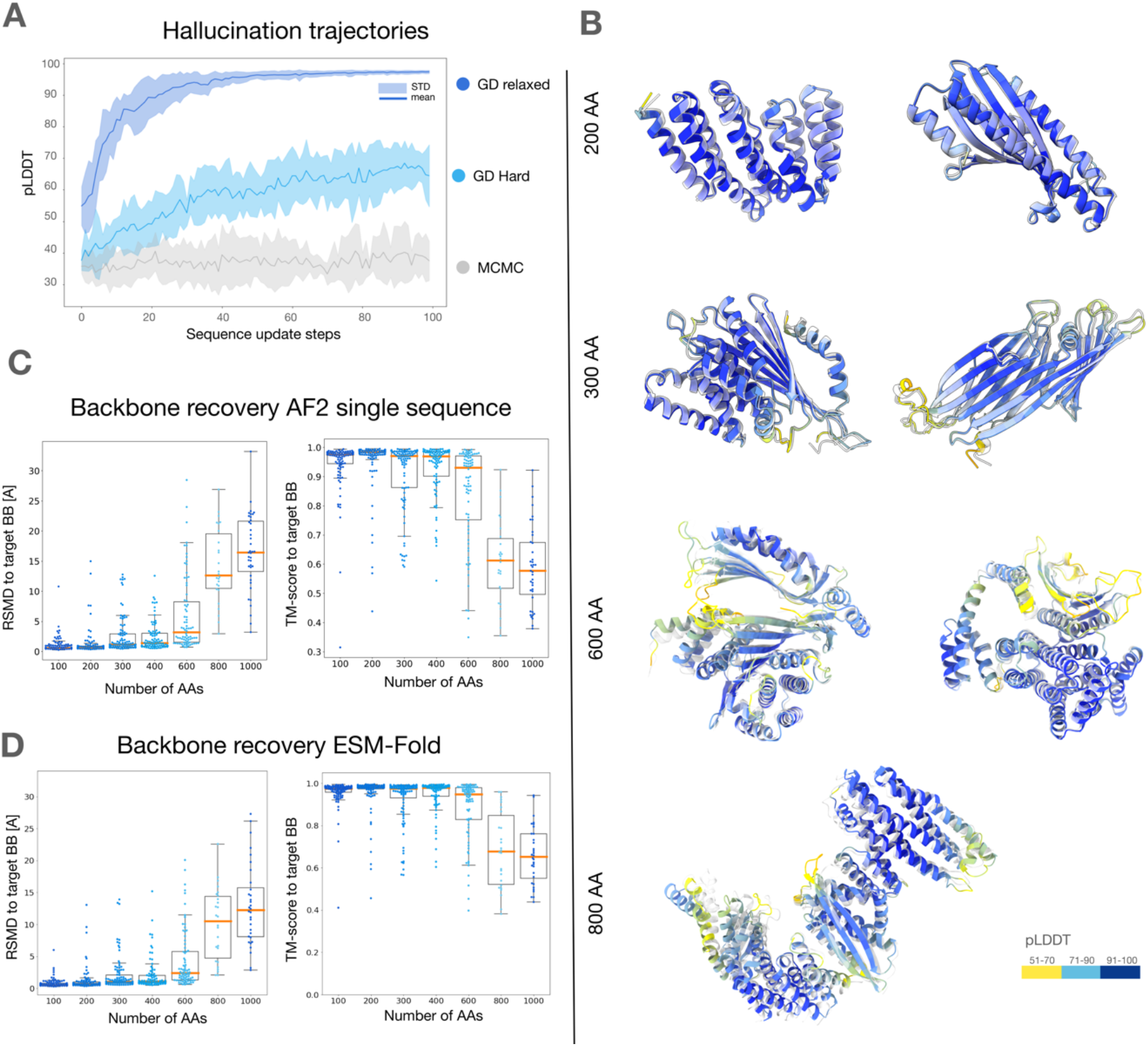
Benchmarking relaxed sequence backbone hallucination. A) Development of the mean pLDDT of 10 design trajectories for unconditional monomer hallucination using relaxed sequence GD, GD with physical sequences (‘hard’) and a MCMC random mutation protocol B) AF2 prediction of unconditional monomers with various length. Grey is the target backbone, while the colored backbone represents the AF2 prediction. Colors indicate pLDDT. C) Backbone recovery of ProteinMPNN designed sequences for unconditional monomers. RMSD and TM-score to target BB in single sequence only mode for 25 −100 hallucinations per size. D) Same as in C) but ESM-Fold was used to predict the structure

The output of GD relaxed backbone hallucination is structurally diverse, including all helical to all beta and combinations of helical & sheet mixtures (Fig.: 2, B) and can also be utilized to hallucinate backbones with multiple subunits. To further benchmark our hallucination approach, we examined the backbone recovery^17^ over multiple sizes of proteins ranging from 100 to 1000 amino acids (AAs). To this end we generated eight candidate sequences for each GD relaxed generated target backbone using ProteinMPNN, and then re-predicted these candidate sequences with AlphaFold2 in single sequence mode. We calculated the root-mean-square-deviation and TM-Score^18^ of the predicted structure relative to the initial backbone input and chose the best hit. We observed high agreement of predictions of MPNN-redesigned sequences to the target backbones over all protein sizes tested (Fig.: 2, C). Judging by the obtained median RMSD relative to the target backbone, relaxed sequence hallucination delivered median sub 4 angstrom backbone recovery for target proteins up to 600 AAs in size. The backbone recovery efficiency drops for backbones larger than 600 AAs (Fig.: 2, C).

To address the potential concern that the generated backbones are adversarial and specific to AlphaFold, we used an independently trained model ESM-fold, which was recently used to generate experimentally validated *de novo* designs^9^, as an orthogonal test. We see ESM-Fold is able to recapitulate the ProteinMPNN redesigned backbones in a similar fashion than AF2 with single sequence input (Fig.: 2, D).

### Experimental validation

To test whether our GD relaxed hallucination protocol could produce real-world expressible proteins we generated a test panel of monomeric protein backbones covering sizes ranging from 100 to 600 amino acids. The geometries were chosen to resemble a mixture of alpha-helical and beta-sheet structural elements. We then fed the resulting backbones to ProteinMPNN for candidate sequence generation and performed AF2 single sequence prediction verification. The candidate sequences were gene-synthesized, expressed in *Escherichia coli* (BL21), and purified through immobilized metal affinity chromatography (IMAC). 13 out of 14 tested protein designs, no matter their size, expressed well, were soluble and had the correct molecular weight as determined by SDS page analysis (Supp. Fig.: 2, A). Size exclusion chromatography elution profiles obtained under native conditions (Fig.: 3 (SEC)) were consistent with the expected sizes of the tested proteins and predominantly showed one single elution peak. All of the SEC tested proteins produced circular dichroism (CD) spectra that indicated that the proteins are well-folded and featured the secondary structure elements that were expected based on their design. All of the tested proteins were also markedly thermostable, as indicated by the CD spectra remaining largely unaltered up to temperatures of 95°C (Fig.: 3, CD). We attribute these satisfactory results to favorable backbone and sequence generation by GD relaxed and ProteinMPNN working in concert and the AF2 single sequence filter. Indeed, one-hot-encoding the relaxed sequence representation *without* passing backbones through ProteinMPNN, resulted in largely insoluble proteins in our hands (Supp. Fig.: 2, B), consistent with previous findings^9^. With a pass through ProteinMPNN, the designed sequences expressed and folded according to expectation (Supp. Fig.: 2, B), in support of previous findings^6,15^. This shows the high value of ProteinMPNN in the design pipeline. Complementarily, the GD relaxed approach now efficiently generates high quality backbones.

**Figure 3.**
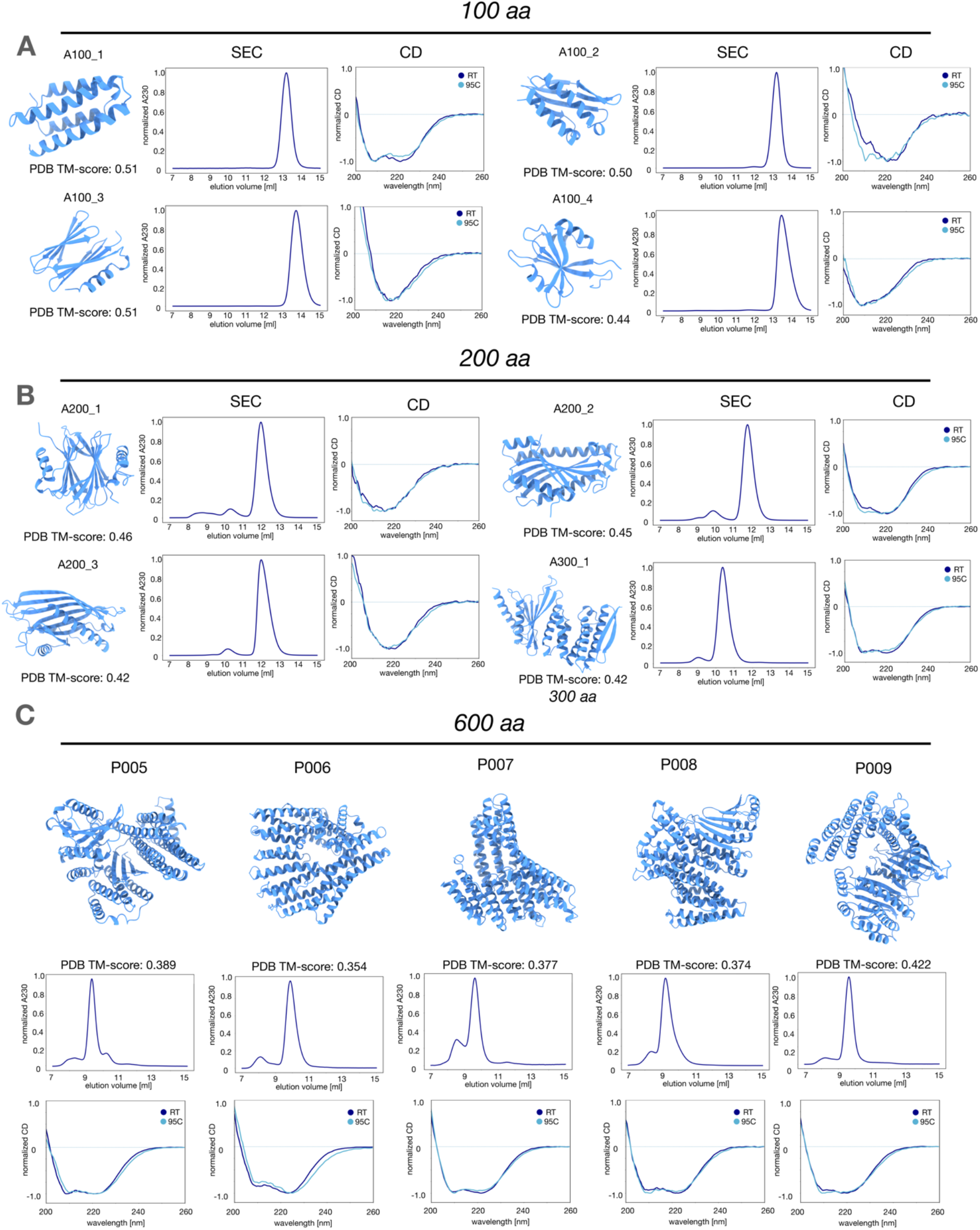
Experimental characterization of monomers. **Models**: AF2 predictions of designed proteins. Cartoon view is created using ChimeraX **SEC**: Superdex Increase 75 10/300 traces of designed proteins. Absorbance at 230 nm is measured to ensure signal to noise ratio. **CD**: Normalized Circular Dichroism spectra of proteins at RT and 95 C. **TM-score:**TM-score is obtained using foldseek^27^ against the PDB100 2201222 A) 4/4 unconditional monomers with 100 AA in size tested. B) 3/3 unconditional monomers with 200 and 1/2 unconditional monomers with 300 AA in size tested C) 5/5 unconditional monomers with 600 AA in size tested

We adapted our pipeline to design multimeric proteins (See Materials & Methods) and tested homo-dimeric and -trimeric candidate designs. All oligomeric designs produced soluble proteins with the expected size of the monomeric chain as determined by SDS-PAGE (Supp. Fig.: 3, A). The oligomeric proteins also had CD spectra that were in good agreement with the expected secondary structure (Supp. Fig.: 3, B). We used FPLC-SEC to evaluate whether the proteins actually formed oligomers as designed. This analysis indicated that out of our design candidates, two homodimers and two homotrimers formed (Fig.: 4, A & B). Next, we modified our pipeline to produce heterodimeric proteins. To this end, we installed a gap of 50 amino acids^19^ in the relaxed sequence input to AF2, which modifies the relative positional encoding to the maximum constant, since the relative encoding is clipped at sequence separation of 32. We hallucinated 1000 backbones and generated three candidate sequences for each backbone design with ProteinMPNN. We then used AF2 single sequence prediction to compute separately the structures of the two single chains in each heterodimers. We also predicted all possible homodimers and heterodimers combinations (Fig.: 4, C) to rank the sequences that form with high confidence heterodimers and have low tendency to form homodimers. We scored the designs based on pLDDT and an interface predicted align error (iPAE) (Fig.: 4, D) that is obtained by summing only over the PAE between the two chains. We experimentally tested 18 design pairs, out of which we selected three candidate design pairs because of superior expression yields. The design pair C5-C6 (Fig.: 4, E) showed two distinct monomer peaks in the SEC when expressed and purified individually. A new peak at larger molecular weight emerged in the elution profile once the purified proteins were mixed. The design pair D9-D10 also showed the designed heterodimer formation when we mixed both chains (Fig.: 4, F). Interestingly, one of the monomer chains (D9) formed a homo-oligomer that disassembled and got replaced with the hetero-dimer when D9 was mixed with D10. AF2 predicts that D9 by itself forms a homo-tetramer. These tests of homo- and heterooligomer designs, while not exhaustive, suggest that our relaxed sequencing hallucination method is also capable of designing functional protein-protein interfaces that specifically interact with each other.

**Figure 4.**
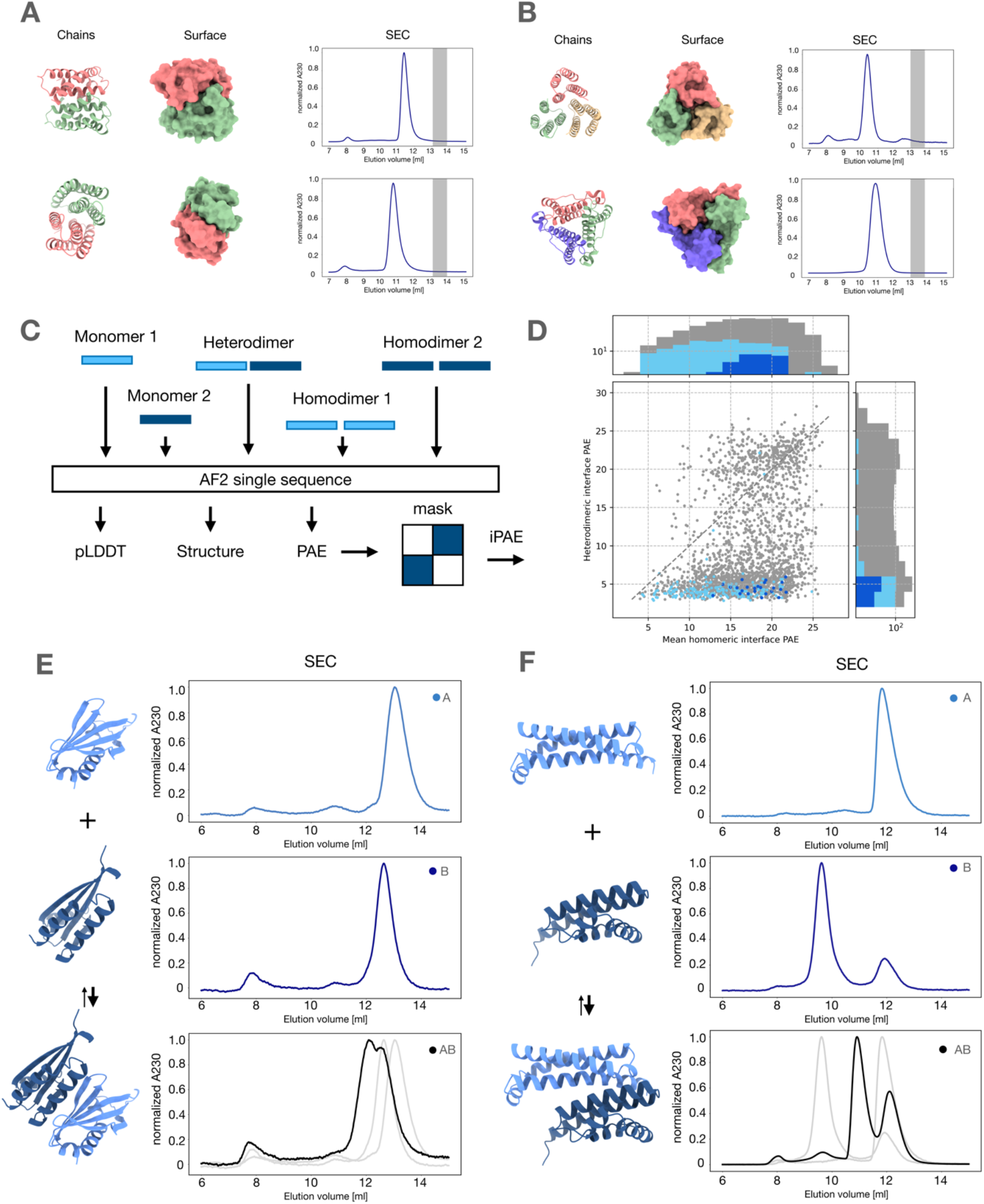
Design and characterisation of homo- and heterooligomers. A) AF2 single sequence predictions of designed homodimers. Solid lines give SEC elution profiles obtained for homo oligomers. The grey area is where one would expect monomer elution based on sizes in Fig 3. B) AF2 single sequence predictions of designed homotrimers. Solid lines give SEC elution profiles obtained for homo oligomers. The grey area is where one would expect monomer elution based on sizes in Fig 3. C) Schematic representation of the heterodimer design pipeline. The colored bars indicate sequences for protomer A & B. All possible combinations are predicted with AF2 single sequence to obtain in silicon benchmarking D) Scatter plot of the i_PAE of the designed heterodimers. Dark blue dots represent designs chosen for experimental testing E) Models and SEC traces of individual protomers and assembled heterodimer pair C5 & C6 F) Models and SEC traces of individual protomers and assembled heterodimer pair D9 & D10. In the combined plot A is added in 1:1.2 excess.

Finally, we validated experimentally the structures of a set of candidate designs using single particle cryo electron microscopy (cryo-EM) analysis. To this end we randomly chose three out of the five proteins comprising 600 amino acids described in Figure 3 with a molecular weight around 60 kDa. We determined 3D cryo-EM maps for all three candidate designs using image data acquired with a Titan Krios cryo EM followed by image processing with RELION^20^. The resulting maps had resolution between 5.7 and 5.9 A according to the FSC gold standard criterion^21^ (Supp. Fig.: 4). At this resolution, the global shape and secondary structure elements can be clearly discerned (Fig.: 5, A, B, C). Rigid body docking using ChimeraX^22^ of AF2 predictions to the measured cryo EM density maps showed good agreement with the designed models. While the resolution was not high enough to obtain atomic models from the densities, using PHENIX real space refinement^23^ we could fit the AF2 predictions to the cryoEM densities and obtain RMSDs of 0.36 A (P005), 0.44 A (P008) and 0.544 A (P009), respectively, between the PHENIX-fitted atomic models and the AF2 predictions. While these fits may not be sufficiently accurate to evaluate details such as side chain geometries, they do indicate that the relaxed sequence pipeline delivers correct output backbone geometries.

**Figure 5.**
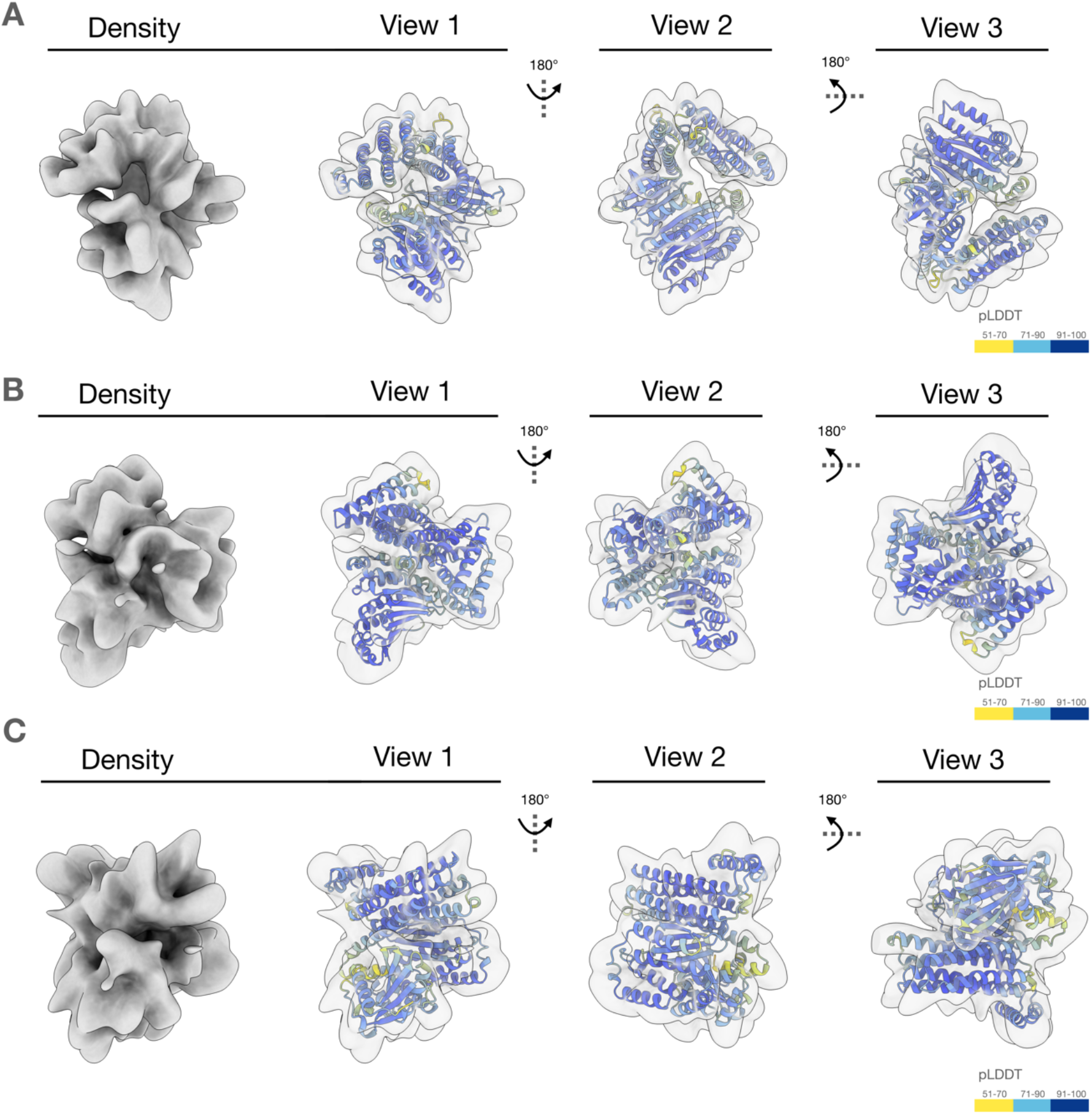
Single particle Cryo EM validation of 600 AA proteins. A) Isodensity surface renderings of the unsharpened 3D cryo EM density of proteins P009 (see SEC and CD Data in Fig. 3) with a final resolution of 5.9 A C). View 1 to View 3 show overlays of AF2 predictions that have been docked using rigid body docking into the densities. B, C) As in A) but for proteins P008 and P005, respectively.

## Conclusion

In this work we introduced and experimentally tested a novel backbone generation method using relaxed sequence hallucination. By enabling the unconstrained optimization of sequence gradients using this relaxed sequence representation, our method can efficiently and accurately generate protein backbones up to 600 amino acids long and possibly beyond. We demonstrated experimentally that the designs translate to real proteins that fold well and have the designed properties. Our approach can generate monomeric proteins but also homo oligomers and heterodimers. The relaxed sequence hallucination method provides substantial efficiency advantages relative to the more commonly used MCMC methods. For example, recently described de-novo designed luciferase enzymes were produced by MCMC hallucination with 30,000 iterations^4^ or large language model based designs requiring up to 170.000 iterations^9^. By contrast, our gradient-descent approach typically converged within < 100 iterations. We thus expect our method to drastically speed up the throughput and scope of protein design tasks. Finally, our relaxed sequence gradient descent provides an alternative to recently introduced denoising diffusion models^17,24–26^, providing researchers with even more tools to enhance the capabilities of de novo protein design.

## Author contribution

C.F. and H.D. designed the research. H.D. supervised the research. C.F. designed proteins and performed computational studies. D.S and C.F. designed heterodimers. Y.d.S designed AF only proteins. S.O. and S.F. developed the ColabDesign Framework. S.O. helped conduct computational studies. C.F. Y.d.S. and D.S. performed wet lab experiments. A.K. prepared Cryo-EM grids, collected data and performed data processing. C.F., A.K. and H.D wrote the manuscript. Y.d.S. and S.O. proofread the manuscript.

## Data & Code availability

All experimental data, sequences, AF2 predictions and Cryo-EM maps will be made available after publication. The exact code used in this manuscript will also be available after publication. The code for the Colabdesign framework is freely available at:

https://github.com/sokrypton/ColabDesign

## Acknowledgement

We thank Google Cloud Services for providing computational resources. We thank Massimo Kube for help with compute cluster maintenance and useful discussions and Maximilian Honemann for guidance with cloning. We thank Justas Dauparas for help debugging the ColabDesign code.

This work was supported by a European Research Council Advanced Grant to H.D. (grant agreement 101018465), the Deutsche Forschungsgemeinschaft through grants provided within the Gottfried Wilhelm Leibniz Program (to H.D.). We (H.D. & C.F.) acknowledge additional support via the Excellence Strategy of the Federal Government and the Länder through the TUM Innovation Network RISE grant. S.O. is supported by NIH DP5OD026389, NSF MCB2032259 and Amgen.

## Supplementary Figures

**Supp. Figure 1.**
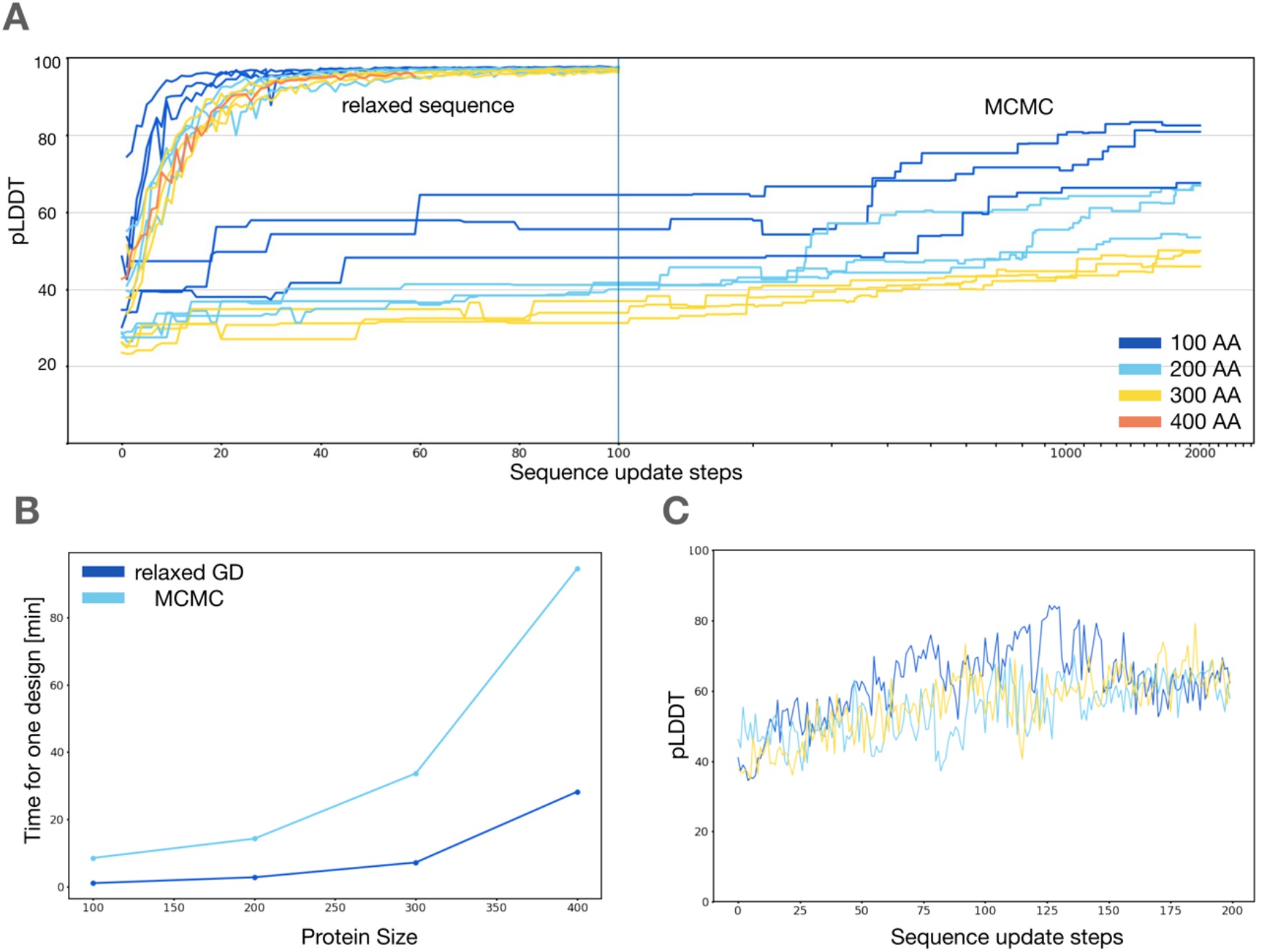
Computational benchmarking. A) Comparison of design trajectories between relaxed-sequence hallucination and MCMC based design. X-axis is linear scaled up until 100 steps and then logarithmic. Both relaxed sequence and MCMC was performed using the ColabDesign framework. A mutation rate of 3 was used for MCMC. B) Time measured to perform one design trajectory using the CD framework on an Nvidia A100. For logits hallucination 100 iterations were used while for MCMC 2000 iterations with a mutation rate of 3 was used. C) Three design trajectories of 200 steps of hallucination performed with the ‘hard’ protocol using ColabDesign.

**Supp. Figure 2.**
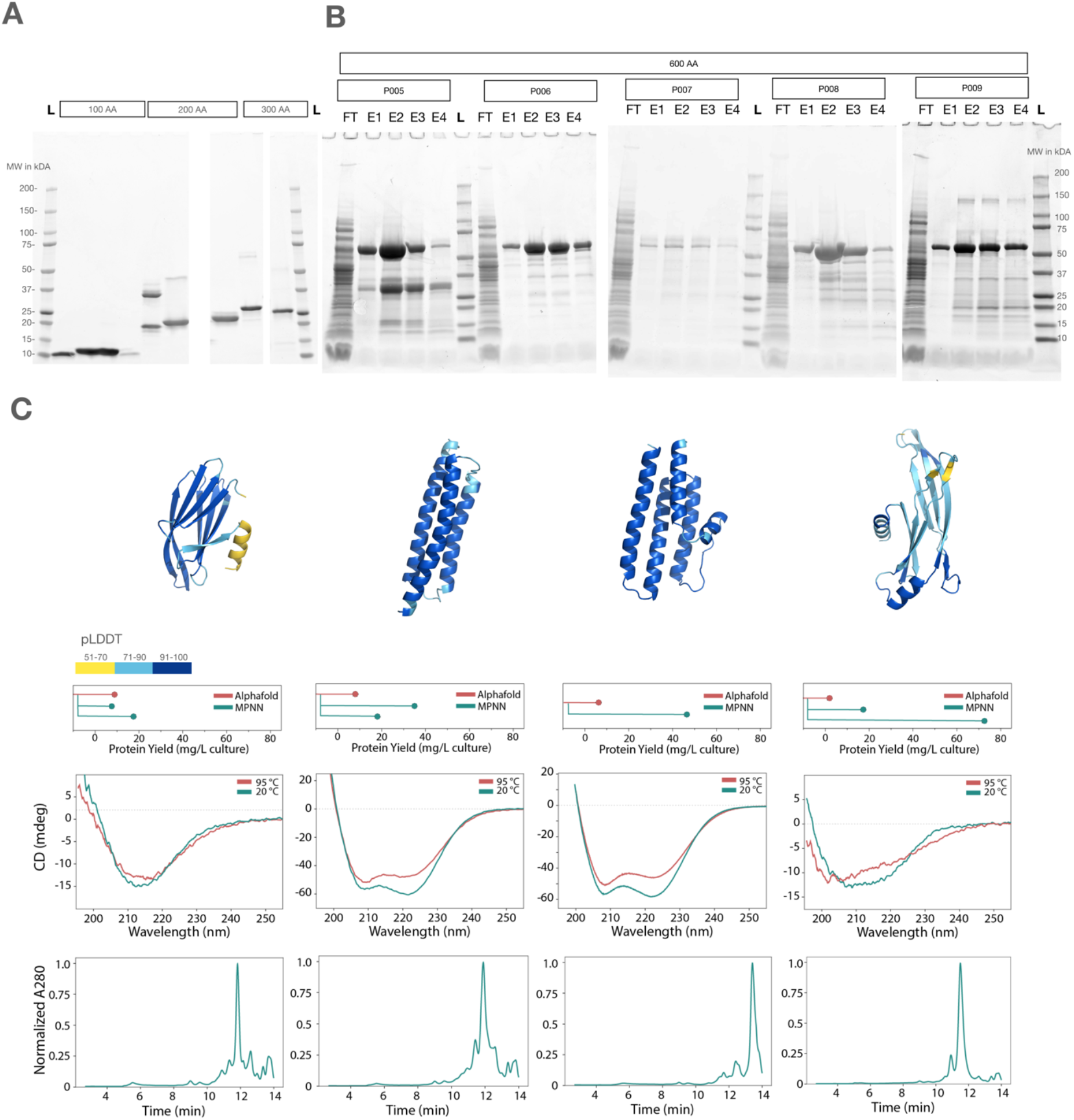
Additional experimental verification. A) Laser scanned images of SDS-PAGE gels analyzing unconditional monomers with 100, 200 and 300 AAs. L indicates the ladder, while the sample bands are elution fractions after IMAC. B) Laser scanned images of SDS-PAGE gels analyzing unconditional monomers with 600 AAs size. E1-E4 indicate the elution fractions after IMAC, while FT shows the flow through C) Experimental testing of AF only generated sequences. TOP: Comparison of expression yield. Yield of protein expression is determined as amount of soluble proteins measured with nanodrop after expression and IMAC purification. MIDDLE: CD spectra of MPNN redesigned backbones. BOTTOM: HPLC SEC traces of proteins indicating their monomeric status.

**Supp. Figure 3.**
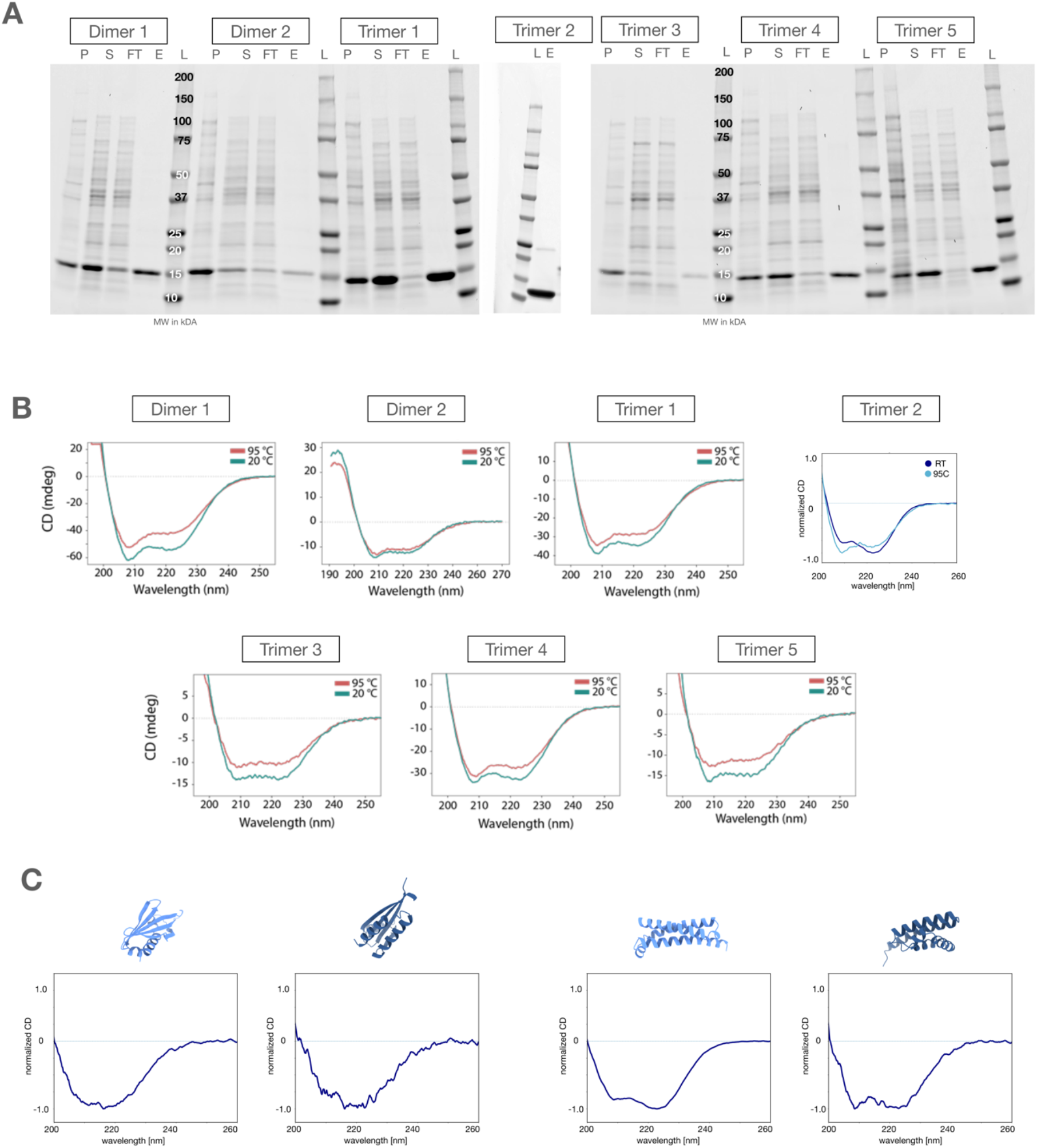
SDS-PAGE gels of Homo Oligomers and CD-spectra. A) Laser scanned images of SDS-PAGE gels showing experimental success of homo oligomer expression and IMAC purification. P = Pellet; S = Supernatant; FT = IMAC Flow Through; E = IMAC elusion fraction; L = Ladder B) CD spectra of homo oligomers C) CD-Spectra of heterodimer protomers measured only at room temperature

**Supp. Figure 4.**
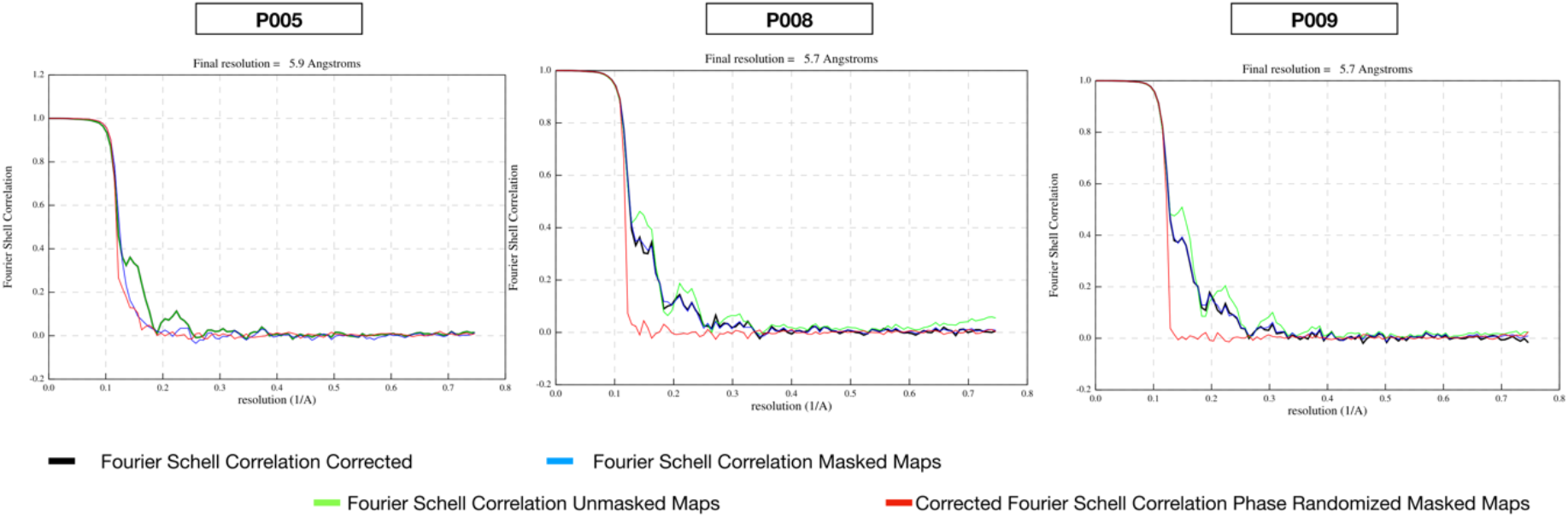
FSC curves of CryoEM densities. FSC Curves created using RELION 4.1 for cryoEM reconstructions shown in Fig.: 5.

## Materials & Methods

### Relaxed sequence hallucination

Hallucinations were performed using the ColabDesign framework either on Google Colab, Google Cloud or on a local installation. Relaxed sequence hallucination was done using the “design_logits” protocol. Softmax of the Gumbel distribution was used to initialize the sequences. Normalized gradient descent was used^28^. The standard loss function consistent of a pLDDT, PAE, contact and radius of gyration losses. Hallucination trajectories were run for 100 iterations. Relaxed sequence hallucination outputs were saved as pdb files with a placeholder sequence generated by taking the argmax of the sequence logits.

### Multimer design

Homo-oligomers were designed using the ‘copies’ option in ColabDesign. This copies the input logits and introduces an index shift of 50 at the breakpoint between the copies. Additionally, the loss function was upgraded to feature interface PAE and contact losses.

### Protein MPNN sequence design

To redesign sequences used for experimental testing a local installation of ProteinMPNN^15^ was used. Batch_size was set to 1 and a sampling temperature of 0.1 was used. Cysteines were omitted to reduce problems in expression. For computational benchmarking a custom JAX-implementation of ProteinMPNN was used (part of the ColabDesign Framework). Again, sampling temperature was 0.1 and the batch_size was set to 8.

### Gene construction and protein expression

Synthetic DNA sequences encoding for the proteins were ordered from IDT as eblock or gblocks and cloned into a pet28b(+) vector using Gibson Assembly (NEBuilder, New England Biolabs). All constructs included a N-terminal His Tag and Heterodimers and 600mers additionally had a c-terminal strep tag. Constructs were cloned into E.coli (DH5a) and positive clones were isolated, miniprepped and plasmids were sequence verified. Verified plasmids were transformed into E.coli BL21 and expressed from a single colony overnight using a homemade autoinduction media (For 1 L: 1x TB media (Carl Roth), 2g lactose, 0.5g glucose). Expression was done in a shaking incubator at 37C in Thomson Ultra Yield flasks.

### Protein purification

Overnight cultures were transferred to 50ml Falcon tubes and centrifuged for 15 min at 4600g to pellet bacteria. Supernatant was removed and the pellet was resuspended in B-Per bacteria lysis reagent (Thermo Fisher scientific) supplemented with Pierce Protease inhibitor (Thermo Fisher scientific). The lysis reaction was carried out for 15min at room temperature with gentle shaking. Lysed cells were pelleted for 10 min at 20.000g. The supernatant was collected and applied to a equilibrated immobilized metal affinity gravity flow column. Column equilibration was done by two column volumes (CV) water and 2 CV wash buffer (1x PBS supplemented with 20 mM imidazole). Flow through was applied a second time to increase yield. The column was washed with 10 CV wash buffer and then eluted using four 0.5 CV elution steps with Elution buffers 1-4 (all four 1x PBS and 100, 200, 300 and 500 mM imidazole). All fractions were analyzed using SDS-PAGE. Eluted proteins were desalted using either Amicon Ultra centrifugation filters (Merc Millipore) or Zeba Spin desalting columns (Thermo Fisher scientific) and concentrations were obtained using absorbance at 280nm.

### CD-spectra measurements

CD-spectra were obtained using a Jasco-815 spectrophotometer. A cuvette with 1mm pathlength was used. Proteins were buffer exchanged into ddH20 and measured at a concentration of 0.3 mg/ml. Measurements at 95C were done by heating up the instrument and samples were allowed to incubate at 95C for 5 minutes before measurement.

### Size exclusion chromatography (SEC)

HPLC-SEC was performed by using a HPLC equipped with a Yarra 3000 HPLC column at a flow rate of 1ml/min and 90-94 bar pressure. Phosphate buffered Saline (PBS) was used as liquid phase. FPLC-SEC was performed on a Äkta Pure 25 using a Superdex 75 Increase 100/300 column with a flow rate of 0.5ml/min at 4C. Again, PBS was used as liquid phase.

### Preparation of Vitrified Specimens

Cryo-electron microscopy (cryoEM) grids, specifically Quantifoil 200-mesh copper grids with R1.2/1.3 holey carbon support films, were first glow discharged for 90 seconds using high-pressure air. The sample was subsequently applied onto the grid within the Vitrobot Mark IV chamber (FEI). Prior to sample application, the chamber was adjusted to 100% humidity at 4°C. After the sample was applied, excess solution was blotted for 3 seconds using a blot force of 20, following which the grid was immediately plunged into liquid ethane for vitrification.

### Data acquisition

Cryo-electron microscopy data was acquired from three different samples, P005 (2800 movies), P008 (3356 movies), and P009 (4211 movies), using a Titan Krios microscope operated at 300 kV (ThermoFisher Scientific) equipped with a Falcon3 direct electron detector and a CS corrector. Movies were taken in nanoprobe mode with a 50 μm C2 aperture and a 100 μm objective aperture, and magnified pixel size of 0.67 Å. Each movie consisted of 50 frames with a total dose of 50 e-/Å2, and an exposure time of 35 seconds with a dose rate of 0.65 e-/pix/s on the detector. Data collection was automated using EPU (ThermoFisher Scientific).

### Data processing

All the movies collected were subjected to motion correction using motioncor2. CTF estimation was performed using CTFFIND-4.1 software on the non-dose-weighted micrographs. Subsequently, particles were picked using a Laplacian-of-Gaussian automated picking routine on the dose-weighted micrographs. The particles were extracted in RELION 4, using a box size of 220 pixels. A low-pass filter was used to reduce the high-frequency details in the predicted atomic model and smooth out its shape to make the initial model. For samples P005, P008, and P009, 1.6 million, 1.8 million, and 2.4 million picked particles were subjected to rounds of 2D and 3D classification, which resulted in a homogeneous set of projections. This process led to 1.5 million, 1.4 million, and 1.9 million particles being used for the final refined job in RELION for proteins P005, P008, and P009, respectively. Additional rounds of classification and refinement could further enhance the resolution of the final refined jobs. The final consensus refinement produced a structure with resolutions of 5.9 Å, 5.7 Å, and 5.7 Å (FSC = 0.143, gold standard) for samples P005, P008, and P009, respectively.

